## Supplementary material for "Speed-detachment tradeoff and its effect on track bound transport of single motor protein"

### Supporting Information

#### Supplementary Tables

Experimental data on the motor speed and the duration of its attachment with the track was obtained from three studies. In two studies on kinesin motor, its speed was varied by change in temperature [1] and its neck linker length [2]. On the other hand, for dynein, its mutational isoforms were used [3]. The speed and duration of the attachment duration was also obtained for the Wild Type forms of other different motor proteins [1–9]. These experimental data, used in Fig. 1 of the main text , are shown in following tables.

| **Temp. (°C)** | **Motor Speed** ($nm/s$) | **Duration** ($s$) | $T_{1/2}$ ($s$) |
|---|---|---|---|
| 20 | 460 | 1.53 | 1.06 |
| 25 | 660 | 1.11 | 0.77 |
| 30 | 980 | 0.92 | 0.64 |
| 35 | 1270 | 0.8 | 0.55 |
| 40 | 1680 | 0.69 | 0.48 |

Table S1: Speed of the Kinesin protein on the Microtubule tracks and their attachment duration at different temperatures as reported in [1].

| **Neck linker** | **Run length** ($nm$) | **Motor Speed** ($nm/s$) | **Duration** ($s$) | $T_{1/2}$ ($s$) |
|---|---|---|---|---|
| WT | 2015 | 414.12 | 4.87 | 3.37 |
| 0P | 2123 | 285.28 | 7.44 | 5.16 |
| 2P | 1830 | 217.80 | 8.41 | 5.83 |
| 4P | 2092 | 156.44 | 13.37 | 9.27 |
| 6P | 1600 | 113.5 | 14.1 | 9.77 |
| 13P | 1677 | 76.69 | 21.87 | 15.16 |
| 19P | 1323 | 73.62 | 17.98 | 12.46 |
| 26P | 1123 | 61.35 | 18.31 | 12.69 |
| 14GS | 1508 | 39.88 | 37.81 | 26.20 |

Table S2: The speed and attachment duration of the Kinesin isoforms with variable neck linker length on Microtubules as measured in [2]. In this study, the length of the neck linker in the Kinesin was modified by the insertion of poly-Proline (P), Glycine (G) and Serine (S) amino acids.

| **Isoform** | **Run length** ($nm$) | **Motor Speed** ($nm/s$) | **Duration** ($s$) | $T_{1/2}$ ($s$) |
|---|---|---|---|---|
| WT | 809 | 65.54 | 12.34 | 8.56 |
| $AAA1_{E/Q}$ | 618 | 50.62 | 12.21 | 8.46 |
| $AAA1_{K/A}$ | 1618 | 28.93 | 55.93 | 38.77 |
| $AAA3_{E/Q}$ | 854 | 33.45 | 25.53 | 17.7 |
| $AAA3_{K/A}$ | 1146 | 39.32 | 29.14 | 20.2 |

Table S3: Motor speed and attachment duration for the mutational isoforms of the Dynein protein as obtained in [3].

| **Ref.** | **Motor protein** | **Run length** ($nm$) | **Motor Speed** ($nm/s$) | **Duration** ($s$) | $T_{1/2}$ ($s$) |
|---|---|---|---|---|---|
| [2] | Kinesin | 2015 | 414.12 | 4.87 | 3.37 |
| [3] | Dynein | 809 | 65.54 | 12.34 | 8.56 |
| [4] | Dynein | 1900 | 85 | 22.35 | 15.49 |
| [7] | KIF1A, C351 | 840 | 137.70 | 6.1 | 4.23 |
| [5] | Kinesin | 1500 | 389 | 3.86 | 2.67 |
| [6] | Myosin V | 1300 | 500 | 2.6 | 1.80 |
| [7] | KIF1A, K381 | 2000 | 769.23 | 2.6 | 1.80 |
| [1] | Kinesin | 704 | 460 | 1.53 | 1.06 |
| [8] | Myosin VI | 226 | 291 | 0.78 | 0.54 |
| [9] | Kinesin I | 910 | 830 | 1.1 | 0.76 |
| [9] | KIF17 | 560 | 1310 | 0.43 | 0.3 |
| [9] | KIF1A | 550 | 1820 | 0.3 | 0.21 |
| [10] | Myosin V | 1600 | 311 | 5.14 | 3.56 |
| [11] | Myosin V | 1300 | 380 | 3.42 | 2.37 |
| [11] | Myosin V | 800 | 550 | 1.45 | 1.00 |
| [11] | Myosin V | 500 | 440 | 1.13 | 0.78 |
| [12] | Kinesin I | 450 | 109 | 4.13 | 2.86 |

Table S4: Speed and attachment duration for various motor proteins as taken from different experimental studies.
